# Ephaptic coupling enables action potential conduction

**DOI:** 10.1101/2024.11.16.623938

**Authors:** J. Waaben, J.P. Hofgaard, G.L. Pick, C. Loisel, T. Engstrøm, N.H. Holstein-Rathlou, S. Poelzing, M.S. Nielsen

**Author notes:** Corresponding author: Morten Schak Nielsen, PhD, Department of Biomedical Sciences, Blegdamsvej 3, Building 12.5, 2200 Copenhagen N, Denmark. Co-corresponding author: Steven Poelzing, PhD, Fralin Biomedical Research Institute at Virginia Tech-Carilion, Virginia Tech, 2 Riverside Cir, R-2118, Roanoke, VA 24015.

## Abstract

**Background:** Cardiac action potentials are believed to propagate solely by electrotonic transmission via gap junctions. In the alternative theory of ephaptic transmission, changes in extracellular electrical fields and ion concentrations may enable a cardiomyocyte to activate its neighbors. The involvement of ephaptic mechanisms remain highly disputed and has been discounted due to the reported inability of an isolated cardiomyocyte to activate another.

**Methods:** Isolated rat cardiomyocytes were subjected to whole cell patch clamp. Transmission of action potentials was tested in current clamp in isolated cardiomyocytes pushed together and in native cardiomyocyte pairs. Sodium channel activation and inactivation was tested in voltage clamp.

**Results:** Isolated cardiomyocytes were pushed together end-to-end and one stimulated to fire an action potential. In 10 attempts, 2 pairs showed successful transmission of action potentials, showing that ephaptic transmission is possible. In native pairs, subthreshold depolarization was transferred between cardiomyocytes of native electrically coupled pairs, however, only after a delay, which is not be explained from electrotonic theory. Once action potentials were elicited in one cardiomyocyte of a native pair, the neighbor was activated with delays under 2 msec. The activation delay did not correlate with gap junctional coupling, again suggesting the presence of an alternative mechanism. In voltage clamp of native cardiomyocyte pairs, one cardiomyocyte was held at -90 mV while depolarizing pulses were applied to its neighbor. Consistently, activation of sodium channels in the depolarized cell led to activation of sodium channels in the hyperpolarized cell, which fits well with ephaptic but not electrotonic theory. Ephaptic transmission requires a shielded local domain. To separate this, we measured steady state inactivation in native pairs. As expected, all channels inactivated at -40 mV. However, when maintaining a potential of – 40 mV for an extended period, sodium current elicited at 0 mV re-emerged by de-inactivation. The de-inactivation at -40 mV support that a shielded domain exists where potassium currents can hyperpolarize the local potential.

**Conclusions:** We show for the first time that ephaptic transmission is possible, that it is most likely in operation, also in the presence of gap junctional coupling, and finally our study suggests that data obtained in single cells may fail to reproduce key features of the electrophysiology of cardiomyocytes that interact with neighboring cardiomyocytes.

## Introduction

The intercellular propagation of the cardiac action potentials is essential for proper activation of the heart, and two theories have been proposed to explain the underlying mechanism. In electrotonic coupling (EtC) through gap junctions, sodium influx during the upstroke of the action potential drives current through gap junctions that depolarizes neighboring cells to threshold for sodium channel activation, which thereby initiates a propagated action potential. In the alternative, ephaptic coupling (EpC), the actual myocardial architecture is taken into account and involves the generation of an extracellular electrical field. Cardiomyocytes are mechanically and electrically coupled at a highly specialized structure called the intercalated disc [1], where the cells interdigitate in a step like manner. The membranes running transverse to the contractile axis in the plicate regions (fascia adherens) contain structural proteins that allow force transmission, whereas the membranes running in parallel in the interplicate regions contain gap junctions that allow intercellular current flow. The membranes at the plicate regions are tightly spaced by the structural proteins leaving only a narrow cleft between the cells. It is often ignored that the plicate regions also contains the majority of the cardiomyocyte’s sodium channels [2] and that this population of sodium channels is particularly responsive to activation [3]. These are facts of important functional consequence, because opening of sodium channels at the intercalated disc not only depolarizes the active cell, but also removes positive charges from the cleft between the myocytes. The removal of positive charge from this confined space creates a local extracellular negative electrical field that effectively depolarizes the opposing membrane due to the close membrane apposition. If the magnitude of this depolarization reaches the threshold for trans-activation of the sodium channels, the neighboring cardiomyocyte will also initiate a propagated action potential. In short, the electrical field coupling theory states that influx of sodium during an action potential generates a local extracellular field that allows cardiomyocytes to activate their neighbors independent of EtC.

There is good evidence for both EtC and EpC. The classical experiments of Silvio Weidmann demonstrated an electrical length constant, which is considerably longer than the length of the cardiomyocytes in Purkinje fibers [4]; and when Barr, Dewey and Berger demonstrated gap junctional plaques in the myocardium, EtC was a convincing candidate for the propagation of cardiac action potentials. Since then experiments have shown that interventions that reduced gap junctional coupling slow conduction [5-7] and that preventing gap junction uncoupling conversely protects the conduction velocity [8, 9]. Nicholas Sperelakis, however, continued to spearhead the concept of EpC by contesting the foundations on which EtC rested, and by providing mathematical models in support of EpC [10, 11]. Since then, modeling work has emphasized the possibility that EpC can act alone or alongside EtC [12-14]. The mechanisms of EpC were further extended in consideration that ion diffusion in the intercalated disc is significantly more limited relative to the interstitium along the lateral membrane, and contemporary models consider ion accumulation and electrodiffusion in their description of EpC [15-17]. Others have gone so far as to model the electrical, diffusional, and anatomical complexity inherent in the intercalated disc to predict the role of ion channel clustering and apposition of sodium channels may play in EpC [18, 19]. Importantly, mounting experimental evidence starts to support the theoretical findings [20-23] and implies the involvement of EpC in cardiac pathologies [24, 25].

The existence and importance of EpC has been highly debated and the mechanism largely dismissed, mainly because arguments for EpC primarily rely on theoretical considerations and indirect experimental evidence. Furthermore, studies in isolated cardiomyocytes showed that transactivation did not occur when cardiomyocytes were pushed together [26], suggesting that the generated extracellular fields were not sufficiently large to support propagation via EpC. However, these experiments were conducted by pushing cardiomyocytes together in a side-to-side configuration and may thus have failed to bring together the regions with the highest sodium current density, which occurs in plicate regions of the intercalated disc. The aim of this study was to test the hypothesis that isolated cardiomyocyte pairs can propagate an action potential by EpC when pushed together end-to-end, and to provide evidence that a protected domain exists in native cardiomyocytes, where EpC can occur.

## Methods

### Isolation of cardiomyocytes

Sprague Dawley rats were anesthetized with isoflurane 5 % in pure oxygen. Once proper anesthesia was obtained a tracheostomy was performed and the animal ventilated using a 7025 Rodent Ventilator, Ugo Basile. Then the heart was exposed, the aorta canulated and the heart retrogradely perfused by gravity with the following solutions containing in mM: 136 NaCl, 2.68 KCl, 1.47 KH_2_PO_4_, 8.06 Na_2_HPO_4_, 0.9 CaCl_2_, 0.9 MgCl_2_, 6 Glucose, pH 7.4, equilibrated with 100 % O2, 37 °C (3 min); then the same solution without calcium (2 min); 20 NaCl, 120 Potassium Gluconate, 1 MgCl2, 10 HEPES, 10 Glucose, pH 7.4, equilibrated with 100 % O2, 37 °C (2 min); then the same solution with 0.025 CaCl_2_, 133-162 U/mL type 2 collagenase from Worthington (15-20 min). The heart was then dismounted from the setup, the atria removed, and the ventricles cut into ∼2×2×2 mm pieces. The pieces were agitated in a solution containing in mM: 100 K-glutamate, 10 K-aspartate, 25 KCl, 10 KH_2_PO_4_, 2 MgSO_4_, 20 Taurine, 5 Creatine, 20 Glucose, 0.5 EGTA, 5 HEPES, 0.1 % BSA, pH 7.4, room temperature (2 min). The solution was strained and calcium raised to 0.056 mM at time zero, 0.166 mM after 10 minutes and 0.717 mM after 20 minutes. The cardiomyocytes were then kept in this solution until use.

### Patch clamp experiments

Cardiomyocyte solution were transferred to a chamber and allowed to settle for approximately 30 seconds. The chamber was then perfused at 1 ml/min at room temperature with a solution containing either: 136 NaCl, 2.68 KCl, 1.47 KH_2_PO_4_, 8.06 Na_2_HPO_4_, 0.9 CaCl_2_, 0.9 MgCl_2_, 6 Glucose, pH 7.4 (experiments where cells were pushed together); or 136 NaCl, 4 KCl, 0.8 MgCl2, 5 HEPES, 5 MES, 0.9 CaCl_2_, 10 Glucose, pH 7.4 (experiments on native cardiomyocyte pairs).

Patch pipettes were pulled from borosilicate glass capillaries (GC150TF-15, Harvard Apparatus) using a PIP 6 temperature-controlled pipette puller, HEKA, to a final pipette resistance between 2-5 MΩ when filled with a solution containing in mM: 140 K-gluconate, 5 KCl, 5 Pyruvic acid, 1 EGTA, 5 HEPES, 0.353 Ca-gluconate_2_, 9 Na-gluconate, 1.92 MgCl_2_, pH 7.2. Two discontinuous switch amplifiers (SEC-05 LX, NPI electronics) were used for current and voltage clamp of cardiomyocytes in the fast whole cell configuration. Data from the amplifiers were filtered using a low pass filter (13 kHz) and sampled at 30 kHz using Cellworks software, an analog-to-digital converter board (PCI1200, National Instruments) and an INT-10 breakout box (NPI Electronic, Tamm, Germany). For stimulation of action potentials, current injections were controlled by an S48 stimulator, Grass Medical Instruments.

### Data analysis

Data were analyzed using custom written MatLab scripts. For analysis of the time point of signal change, the MatLab standard implementation of the CUSUM method was applied; for delays between action potential upstroke, the maximal dV/dt was determined as the point of activation. Data are presented as mean±SEM. Statistical analysis was applied as described in text.

## Results

### Ephaptic AP transmission in de novo formed cardiomyocyte pairs

Cardiomyocytes have the highest expression of sodium channels at the plicate regions of the intercalated disc [2, 3], regions that are located at the end of the myofibrils. Therefore, we pushed cardiomyocytes together end-to-end, to optimize the conditions for EpC to occur, and more importantly, to mimic the natural cellular anatomy of coupling as close as possible. Each cardiomyocyte was zero-current clamped in whole-cell configuration in order to monitor membrane potential and current injections were applied to elicit action potentials (APs). Initially the cardiomyocytes were moved into close apposition, before the solution was switched to calcium free solution to inhibit any auto-adhesion between plicate membranes of the intercalated discs. After 3-5 minutes, the cardiomyocytes were pushed together, and calcium reapplied. APs were elicited in one of the cardiomyocytes (active), while the other cardiomyocyte remained unstimulated (passive). Figure 1A shows a bright field image of two isolated cardiomyocytes that were pushed together end-to-end and Figure 1B shows the voltages from the actively depolarized cell (red) and passive cell in blue. The stimulus depolarized the active cell, however, no electrical coupling between the cardiomyocytes was observed as evidenced by the stable voltage in the passive cell during the stimulus. The lack of electrical coupling was subsequently confirmed in voltage clamp by applying a negative potential to the active cell and observing no current transfer to the passive cell. Data are shown in Figure 1B insert where the red trace shows the current added to hyperpolarize the active cell from -80 mV to -110 mV. Importantly, no current flowed to the passive cell (flat blue line), which was held at –80mV.

**Figure 1.**
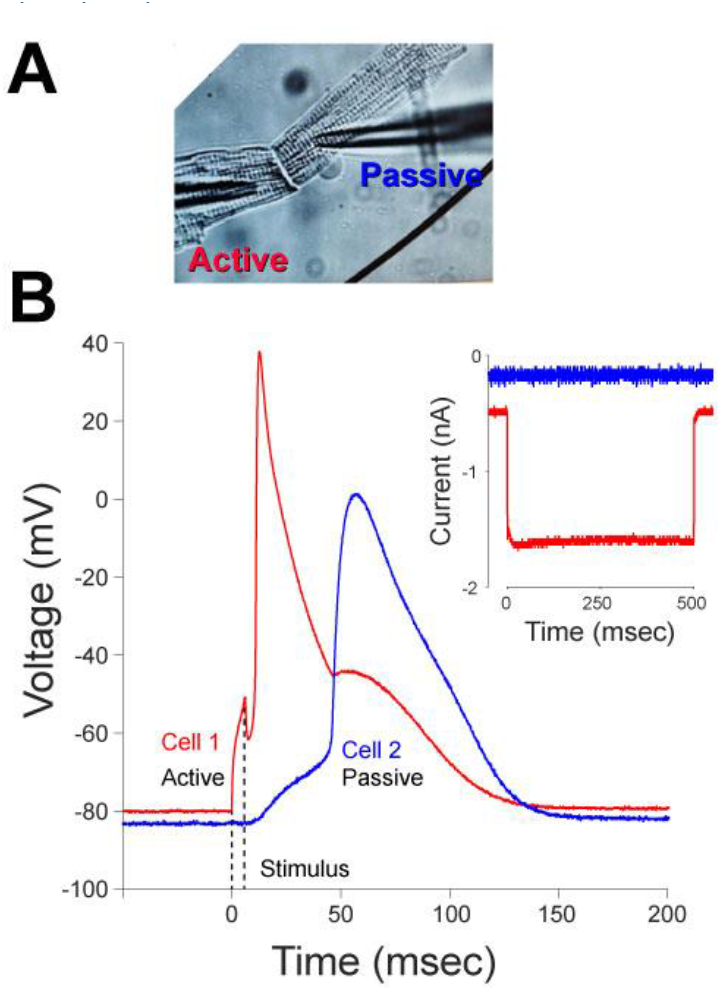
Action potential transmission can occur by ephaptic mechanisms alone. **A)** Bright-field image of two isolated cardiomyocytes that were pushed together end to end. **B)** Action potentials elicited by current injection in the active cardiomyocyte (cell 1, red), stimulus duration indicated by dashed lines. The passive cardiomyocyte (cell 2, blue) subsequently fired an action potential. Insert demonstrates the complete lack of electrical coupling. The active cardiomyocyte was hyperpolarized by current injection (red) and despite the 50 mV electrical gradient between the cells, no current was recorded in the passive cardiomyocyte (blue).

Despite the lack of direct EtC gap junctional coupling, the passive cell started to depolarize once the AP was triggered in the active cell. The membrane de-charging of the passive cell occurred over the time course of the upstroke and continued during early repolarization of the active cell. Consequently, the upstroke phase of the passive cell 2 occurs during repolarization of the active cell. In other words, Figure 1 demonstrates that an action potential can propagate between two isolated cardiomyocytes without measurable gap junctional coupling and therefore that AP propagation can occur by ephaptic mechanisms alone.

Interestingly, the electrical interaction between the cardiomyocytes was bi-directional. Specifically, Figure 1B reveals that the AP of the passive cell was associated with a transient depolarization of the active cell, creating a discontinuity in its repolarization.

The experimental procedure was quite challenging. In the 30 attempts to observe EpC mediated propagation, 20 failed due to lack of success in end-to-end apposition or loss of seal during manipulation. In the remaining 10 experiments, end to end apposition was obtained, but AP transmission was only observed in 2 experiments. The delay between the maximum upstroke of the active and passive cell was 28.2 msec on average. This value is clearly incompatible with the expected propagation time. If the entire delay occurred at the intercalated disc, it would amount to 0.1 msec at a conduction velocity of 1 m/sec and myocyte length of 100 μm.

### Ephaptic AP transmission-Native Cardiomyocyte Pairs

Native cardiomyocyte pairs (Figure 2A) responded differently from the de novo formed pairs. Figure 2B shows that the passive cardiomyocyte (blue) started to depolarize already during the application of the stimulus the active cell (red), which apparently makes a strong case for EtC as the dominant mechanism of propagation in coupled cells. Some features, however, did not support gap junctions as the primary driver for intercellular communication.

**Figure 2.**
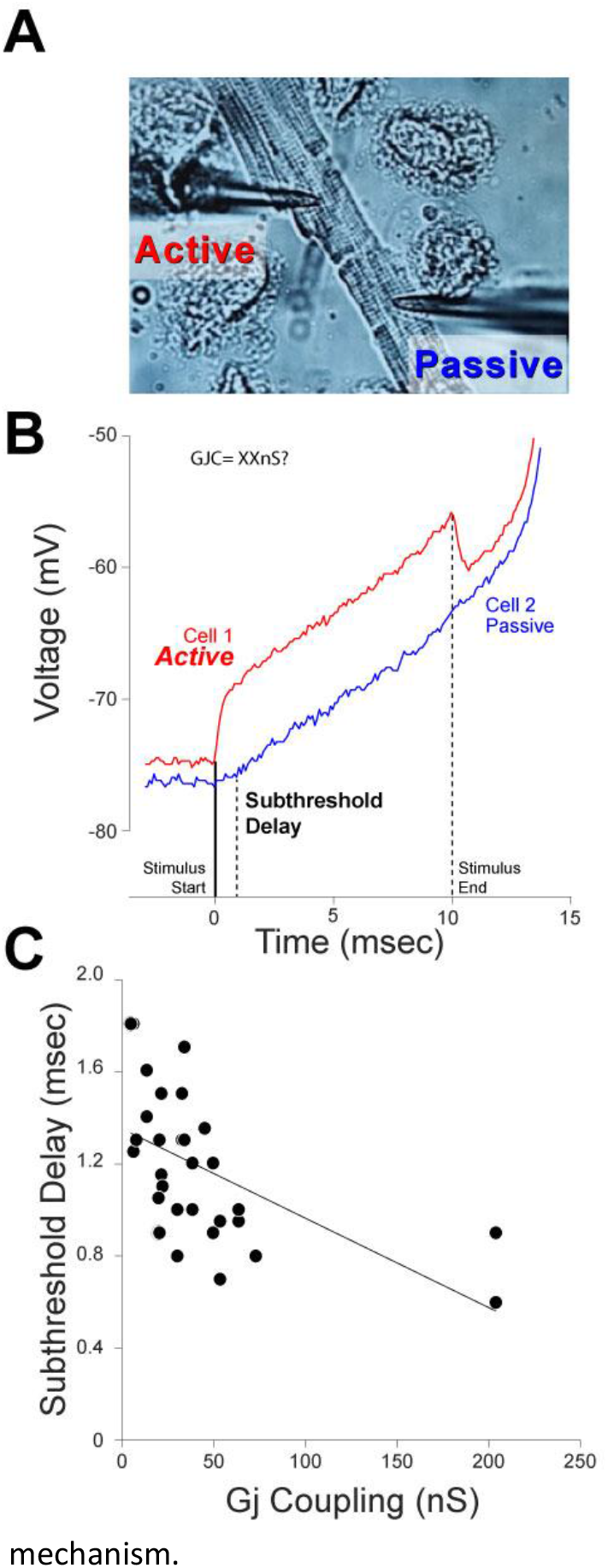
Stimulus transfer between native cardiomyocyte pairs. **A)** Bright-field image of a native cardiomyocyte pair. **B)** Voltage response to a depolarizing current injection in the active cardiomyocyte (cell 1, red) as indicated by the vertical solid lines. Blue trace shows the voltage response of the passive cardiomyocyte, which depolarized with some delay as indicated by the vertical dashed line. The subthreshold onset delay was determined by the CUSUM method. **C)** The subthreshold onset delay of response in the passive cardiomyocytes as function of gap junctional coupling (n=17).

During the stimulus, a closer inspection of the depolarization in the passive cells revealed a considerable delay in the onset of depolarization, which is not expected based on EtC theory. In 17 cell pairs, we determined the gap junctional coupling and time to depolarization onset in the passive cell, which are plotted in Figure 2C. The subthreshold onset delay tended to decrease with increased coupling and a linear fit indicated a weak correlation (R^2 0.37), although the slope was significantly different from zero (p=0.0004). Since gap junctional coupling should be a prime determinant of charge transfer according to EtC, the low predictive power of GJC argues that some other mechanism must be at play. Furthermore, upon termination of the stimulus, the potential of the active cardiomyocyte was markedly reduced and approached that of the passive cardiomyocyte. The resulting drop in driving force across gap junctions had no effect on the slope of depolarization in the passive cardiomyocyte, suggesting that its depolarization is mediated by an intrinsic and non-gap junction mechanism.

Once an AP was elicited in native cardiomyocyte pairs, the upstroke in native isolated pairs was near simultaneously (Figure 3A, left panel, cell 1 active), in contrast to the long delay observed in de novo formed pairs. In the representative example, the pair had an intercellular coupling of 64 nS and the activation delay was 0.1 msec (Figure 3A, insert). However, when stimulating cell 2 of the same pair (Figure 3A, right panel), cell 1 (now passive) still activated first, and consequently the activation delay became negative. This held true for most pairs and Figure 3B shows the activation delay when stimulating cardiomyocyte 2 against the activation delay when stimulating cardiomyocyte 1 (n=37). Most data points lie on or close to the line of inverse identity (slope of -1), meaning that a positive delay going from one cell to another would result in negative delay going in the other direction. In other words, the same cell would always fire first, irrespective of which cell received the current injection.

**Figure 3.**
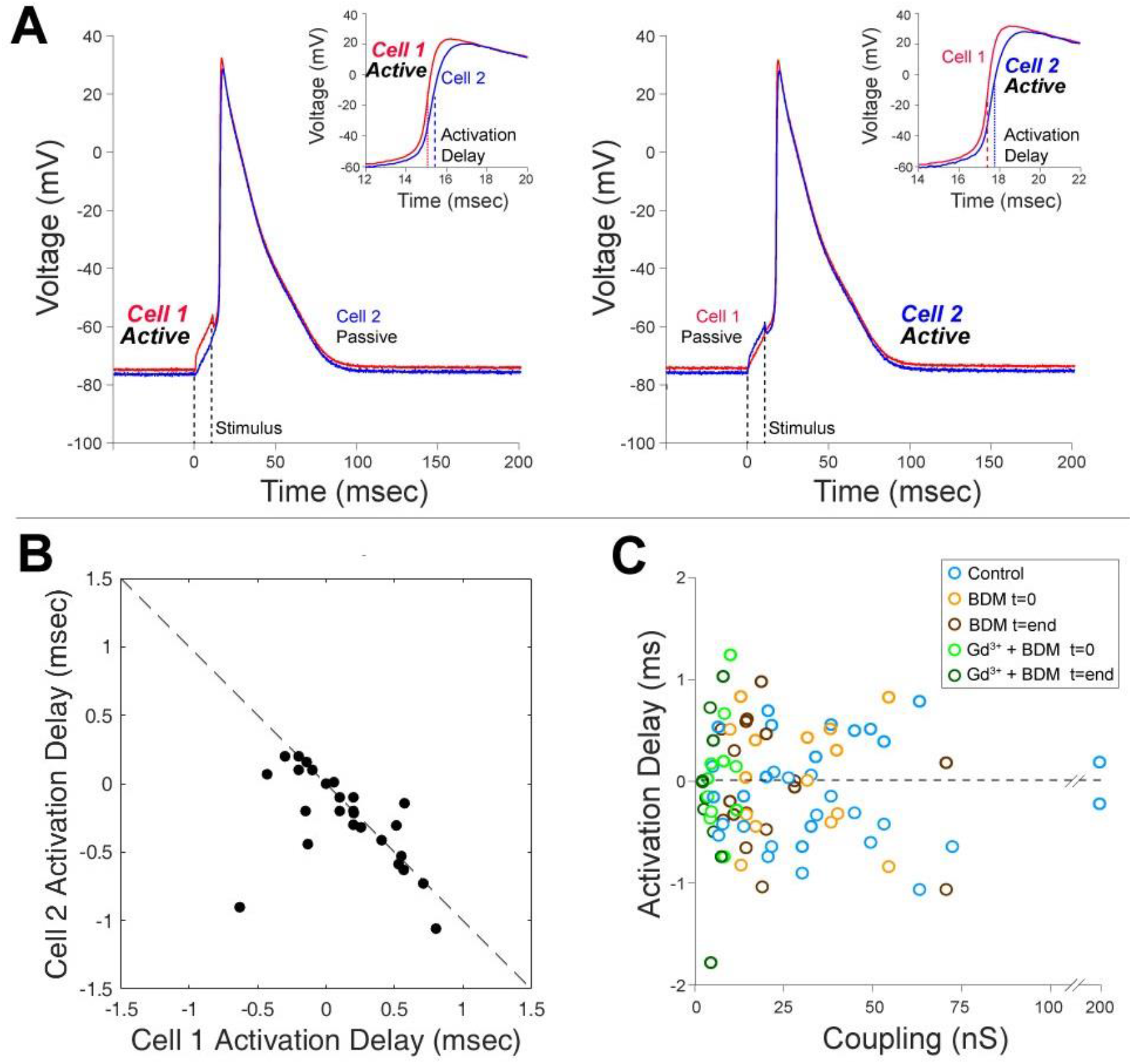
The delay in action potential transmission is determined by cardiomyocyte excitability and is independent of gap junctional coupling. **A)** Left: Action potentials elicited by current injection in the active cardiomyocyte (cell 1, red), stimulus duration indicated by dashed lines. The passive cardiomyocyte (cell 2, blue) subsequently fired an action potential. Insert shows a zoom around the time of activation, showing a short delay between the active and passive cardiomyocyte. Right: Same as left, only cell 2 was actively stimulated. The insert shows that despite cell 2 (blue) being stimulated, cell 1 (red) still fired an action potential first. **B)** Plot of the activation delay when cell 2 was active as function of the delay when cell 1 was active n=37. Dashed line indicates line of inverse identity. **C)** Activation delay as function gap junctional coupling in cell pairs that were either untreated (n=18), BDM treated (n=8) or Gd3^+^ + BDM (n=7). Measurements for treatments were made immediately after obtaining the whole cell configuration and at the end of the experiments

In order to dissect the role of gap junctional coupling in AP transmission, we measured action potential upstroke activation delay and intercellular coupling in untreated native cell pairs, cell pairs exposed to BDM (a commonly used electromechanical uncoupler, which also uncouples gap junctions), or cell pairs exposed to BDM and gadolinium (a gap junction hemi-channel pore blocker) in the bath and pipette to inhibit gap junctional coupling. Data were included in this experimental dataset only if the seals and whole-cell recordings could be obtained at the beginning of the intervention (start) and after some variable period of treatment (end). On average, treatment was 41±7 minutes for BDM and 29±7 minutes for BDM+ gadolinium. Since the aim of these experiments was to induce a wide range of GJ-conductances, the duration of treatment was of no particular relevance as long as gap junctional coupling varied. Although the differences in gap junctional coupling did not reach statistical significance for all groups, the average numerical values fell with the degree of intervention as well as from start to end of each individual intervention.

Figure 3C shows the delay in action potential propagation as a function of GJ coupling. Neither intercept nor slope were significantly different from zero determined by a linear fit to the data, indicating a lack of dependence between activation upstroke delay and GJ conductance.

In summary, the lack of GJ-dependence on AP-delay is at odds with the predictions of EtC, and even at extremely low degrees of coupling, no increase in delay was observed. Furthermore, the fact that activation depended more on the excitability of a given cell, rather than which cell was stimulated, also speaks against a pure EtC mechanism and suggests that somewhere in the cell-to-cell interface, a domain must exist, where transactivation can occur. To further pursue this, we activated sodium channels of cardiomyocyte pairs in voltage clamp mode.

### Sodium channel transactivation in voltage clamp

In theory, it should be possible to prevent EtC dependent trans-activation of sodium channels by clamping a cardiomyocyte of a pair to a hyperpolarized potential while activating the sodium channels in its neighbor. To test this concept, we performed experiments with native cell pairs in voltage clamp, where the active cell was clamped from a holding potential of -90 mV to increasingly depolarized potentials (Figure 4A, left panel), while the passive cell was clamped to a constant potential of -90 mV, a potential that should prohibit the activation of its sodium channels. The representative example in Figure 4A (right panel) shows that a depolarization of -40 mV or more evoked a substantial sodium current in the active cell as expected. The sodium current was quantified as the difference between the minimum value and the steady state current, and the average current is shown as function of voltage in the right panel of Figure 4 (red, n=6). In the passive cell of this and all experiments, sodium channel activation in the active cardiomyocyte led to a simultaneous trans-activation of a sodium current in the passive cell (Figure 4 middle panel, mean values shown in right panel). Gap junctional coupling was calculated from the steady state currents. Trans-activation is not expected from EtC theory and persisted at electrical coupling as low as 18 nS in the example shown (average 57±12.5 nS). It was, however, noted that maximum current in each experiment was seen at the first step sufficient to activate a substantial sodium current. Because of differences in activation thresholds, the average IV curves (Figure 4 left panel) exhibited a gradual increase in current that resembles a normal sodium current IV curve. The ‘all or none’ activation of the sodium current most likely occurs due to inability of the clamp to compensate fast enough once sodium channel activity is initiated. It may be argued that the inability to sufficiently clamp the active cell also prevents the control of the passive cell.

**Figure 4.**
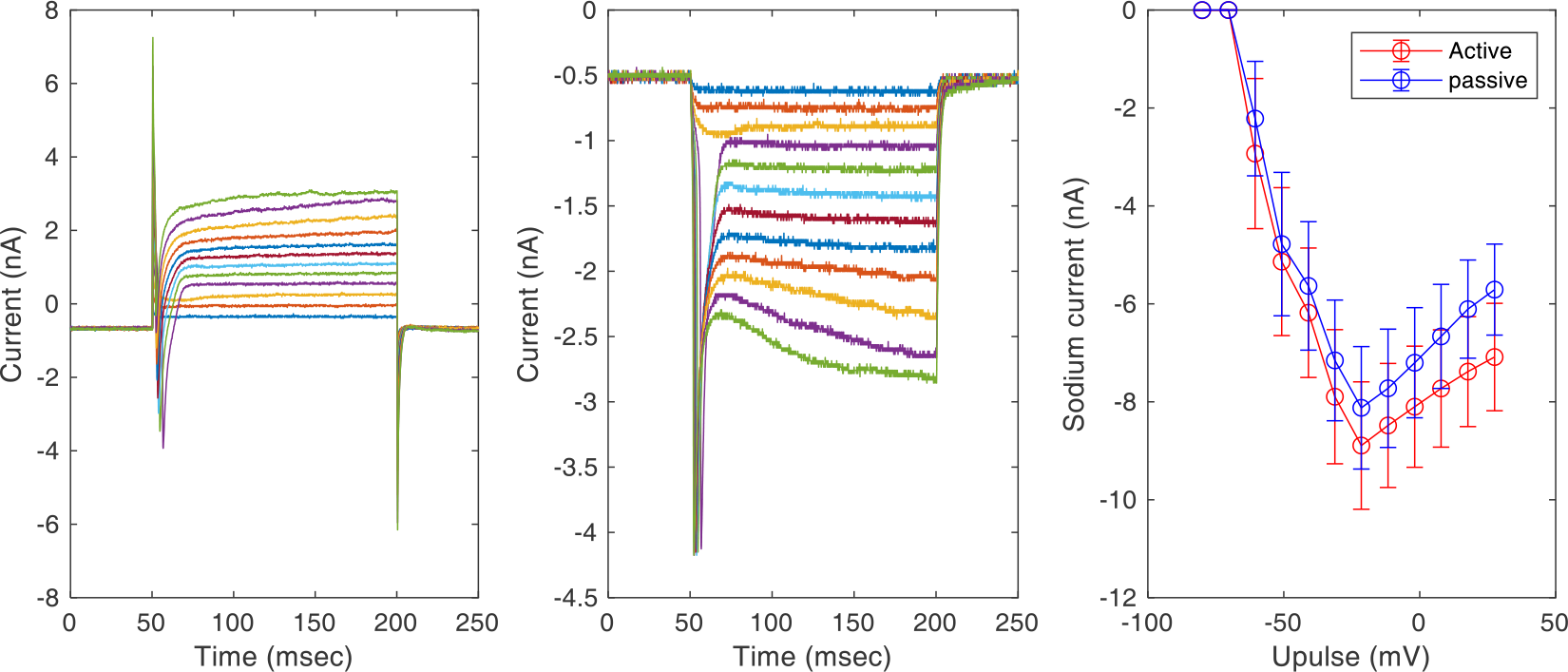
Sodium channel activation in a cardiomyocyte of a pair always triggers sodium channel activation in its neighbor. Left panel: Application of depolarizing pulses of -40 or more positive elicited a sodium current in the active cell. Middle panel: Corresponding current traces from the passive cell. The steady state currents represented the gap junctional current from which the electrical coupling was calculated. Steps that activated sodium channels in the active cell, also stimulated sodium currents in the passive cell as indicate by the large peak negative currents. Right panel: Average sodium currents as a function of voltage step. Currents were determined as the peak negative current minus the current after sodium channel inactivation.

However, given the presented evidence that non-gap junctional mechanisms contribute to AP transmission, another explanation for the trans-activation, could be that shielded domains exist in which the extracellular, and thereby the transmembrane potential, cannot be controlled.

### Providing evidence for a protected EpC domain

A central prerequisite for EpC is the existence of partly shielded domain, where local extracellular potentials and ion concentrations can vary relative to the bulk extracellular space. To substantiate the existence of such a domain and to isolate the sodium currents arising therefrom, we devised a protocol to steady state inactivate the sodium channels facing the bulk bath solution, while allowing those in the isolated domain to de-inactivate. The blue curve in Figure 5B shows the results of a standard steady state inactivation curve where both cardiomyocytes of a pair were held at -120 mV from which 100 msec simultaneous depolarizing pre-pulses were given in steps of 5 mV from -115 to -40 mV in both cardiomyocytes. At the end of the pre-conditioning pulses, the available sodium current was measured by applying a 50 msec test pulse to -40 mV. The protocol results in a sigmoidal curve, where all sodium current is inactivated at -40 mV (n=30 cardiomyocytes from 15 pairs). In our modified protocol, the timing of pulses was identical, but the pre-pulses were varied from -115 to 0 mV and sodium current measured from a 0 mV test pulse. Representative currents are shown in Figure 5A and the average data in 5B (red, n=30 cardiomyocytes from 15 pairs). In contrast to the sigmoidal steady state inactivation observed with a -40 mV protocol, the curve was biphasic with a pronounced foot of currents that could be elicited at pre-pulses positive to -40 mV. Such behavior is incompatible with the normal properties of sodium channels facing bulk solution, and we propose that these currents arise from domains between the myocytes; domains sufficiently shielded that their potassium channels can repolarize the local potential and enable de-inactivation of the sodium channels residing there. To test this assumption, experiments were performed using the 0 mV protocol under control conditions before (Figure 5C, red) and after inhibiting the I_K1_-current by application of 20 μM barium (blue). Barium significantly reduced the foot currents, which we quantified as the area under the curve between -40 and 0 mV (control: 1.34±0.17 vs barium: 0.42±0.08, p<0.001 in paired t-test), demonstrating that repolarizing potassium currents are indeed necessary to enable the paradoxical sodium current that arise, presumably, from the shielded domains between the myocytes.

**Figure 5.**
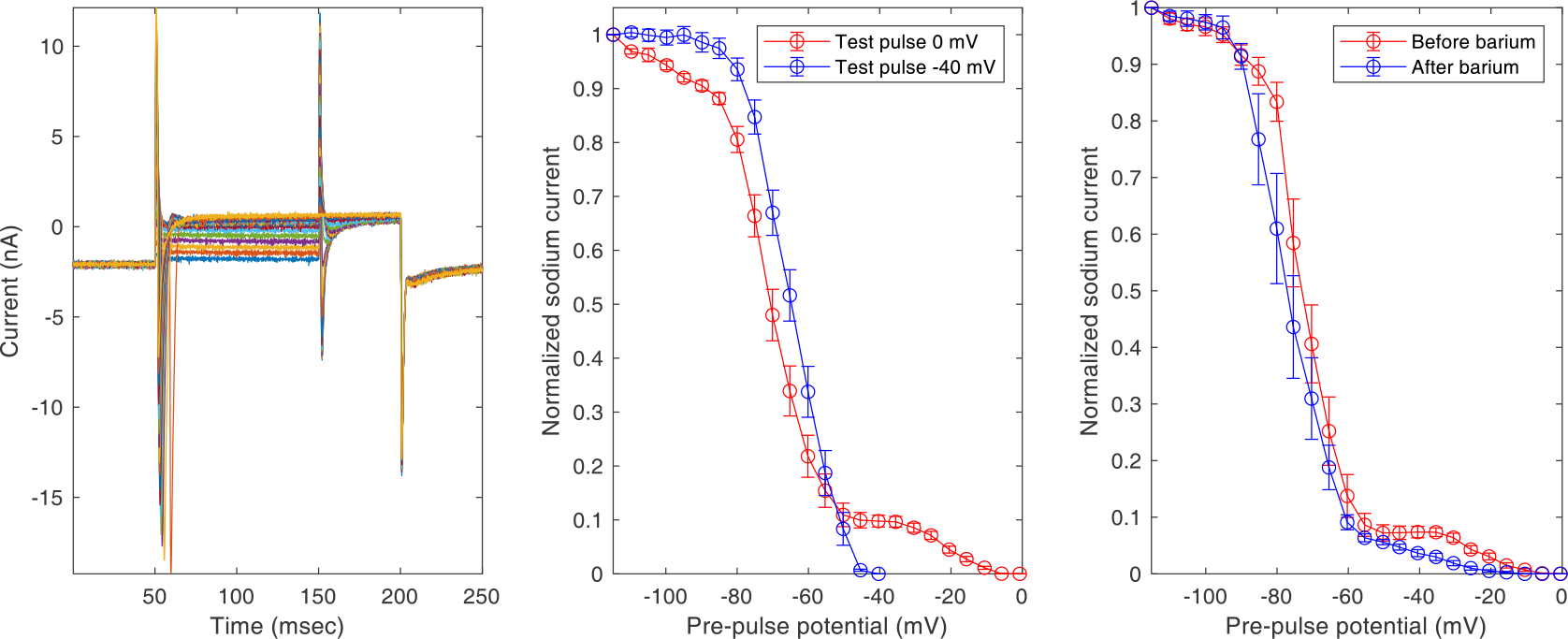
Steady state inactivation of sodium channels. Left panel: Representative example of current traces from an experiment where both cardiomyocytes of a pair were held at -120 mV from which 100 msec depolarizing pulses were applied from -115 to 0 mV in steps of 5 mV to progressively inactivate sodium channels. Finally, a 50 msec pulse to 0 mV was applied to probe the available sodium channels. The same pairs were also exposed to an identical protocol only with a maximum pre- and test-pulse of -40 mV. Middle panel: Sodium current normalized to maximum sodium current available at -120 mV. Data from protocol with 0 mV protocol in red and -40 mV in blue (n=30 cardiomyocytes in 15 pairs). Right panel: Cardiomyocyte pairs subjected to the 0 mV protocol in untreated conditions and after treatment with Ba^2+^ (20 μM).

To further demonstrate that the foot currents arise from de-inactivated sodium channels, we depolarized the cardiomyocytes by a pre-pulse to -40 mV for 5 to 200 msec in 5 msec increments periods before imposing a 0 mV test pulse. The average sodium current is shown as a function of pre-pulse duration in Figure 6. No sodium current was observed at the 5 msec pre-pulse, showing that the foot current channels are indeed inactivated by the pre-pulse. With increasing pre-pulse duration, foot currents increased, and although the curve appears linear, each experiment had a variable time constant of recovery. In 11 experiments, the average time constant was 143±22.8 msec. These data supports that a shielded domain repolarizes and allows the resident sodium channels to de-inactivate.

**Figure 6.**
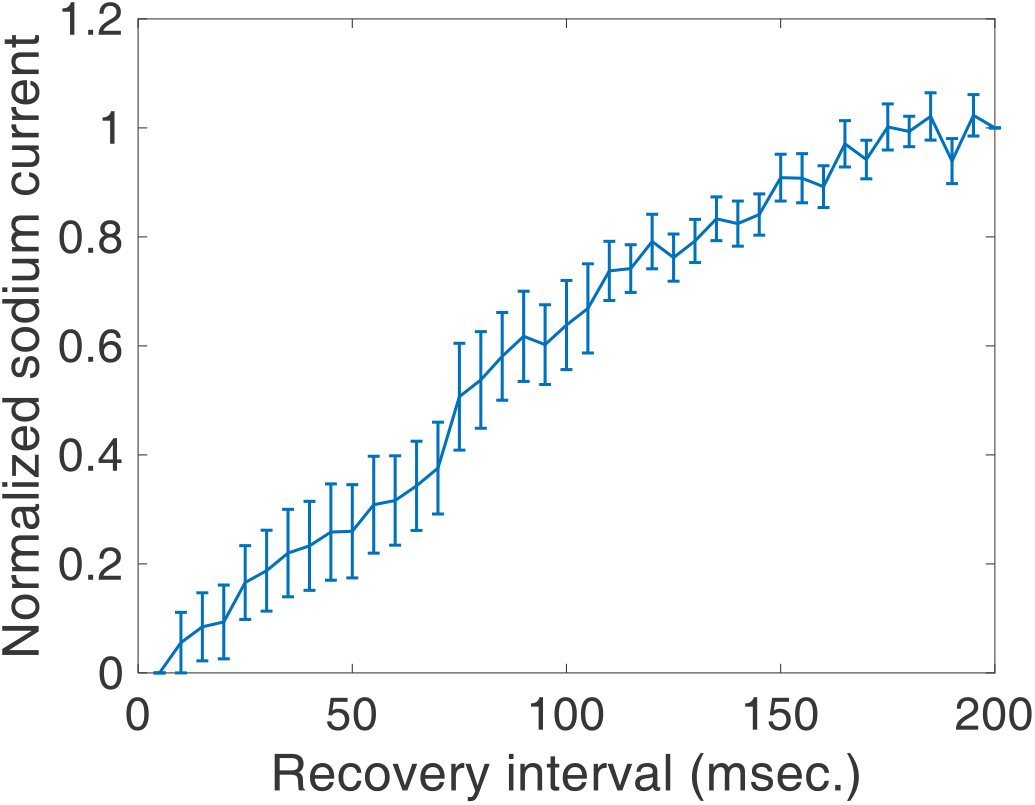
Recovery of sodium channels from steady state inactivation in cardiomyocyte pairs. Both cardiomyocytes of a pair were held at -120 mV from which depolarizing pulses to -40 mV were applied for between 5 and 200 msec. Then, a 50 msec pulse to 0 mV was applied to probe the available sodium current. Graph shows the available sodium current normalized to sodium current available at -120 mV.

## Discussion

In summary, this manuscript demonstrates that cardiomyocytes can activate each other by ephaptic transmission alone, substantiates that ephaptic transmission contributes to AP transmission in native cardiomyocyte pairs and for the first time demonstrates that even isolated cardiomyocyte pairs retain shielded domains, a prerequisite for ephaptic transmission.

The concept of electrical transmission via extracellular fields is not new in biology. At the macroscopic level Carlo Matteucci, showed in 1842 that the exposed heart from one frog can excite the sciatic nerve from a second frog, as demonstrated by the resulting muscle contraction [27]. Nearly a century later, Bernhard Katz and Otto Schmitt demonstrated that stimulating a neuron will hyperpolarize the shared extracellular space sufficiently to activate an adjacent neuron [28]. Shortly thereafter, Arvanitaki proposed the term ephapse to describe the structure, or lack thereof, that supported these phenomena [29]. Meanwhile, strong evidence was provided by pioneering work in electron microscopy and electrophysiology, making a strong case for direct electrical coupling through gap junctions (for review see [1]). However, Nicholas Sperelakis continuously provided evidence that GJ mediated cell-to-cell communication insufficiently explained a number of experimental findings [11, 30-35]. Interestingly, the first demonstration of EpC communication in heart was reported in 1984 by Suenson. He demonstrated that when two papillary muscles were physically tied together, stimulating one muscle would electrically activate and excite the other muscle bundle [20]. Shortly thereafter, Weingart and Maurer pushed two ventricular myocytes together and reported that they failed to transmit APs. Based on this finding they concluded that “*This finding rules out the possibility of an ephaptic mechanism of impulse transmission in cardiac tissue*” [26]. The paper by Weingart and Maurer has often been referenced to discount the possibility of ephaptic transmission, however, this conclusion was based on pushing cells together lateral edge to lateral edge, and at the time the sub-cellular localization of sodium channels on the myocyte was unknown. The reason that we now demonstrate transmission in pairs pushed together in the absence of direct electrical coupling may have several reasons. Probably greatest importance, we ensured that the cardiomyocytes were pushed together end to end, to bring into proximity the intercalated disc plicate regions containing the highest density of sodium channels. Secondly, we removed extracellular calcium before pushing them together. This was done to prevent calcium dependent adhesion molecules like N-cadherin from making bindings between the membranes in the freely exposed plicate region; bindings that could prevent the proper formation of inter-cardiomyocyte adhesion. Lastly, the success rate was very low in our hands and the paper by Weingart and Maurer shows only one example but has no mention of the number of attempts made. In principle, many experiments are needed to substantiate that something is impossible, whereas showing that something is possible only requires one successful demonstration. The delays observed in our two successful demonstrations were clearly incompatible with normal conduction velocities but given the difficulty in obtaining the necessary apposition of sufficient sodium channels across a suitably narrow ephapse, it is likely that conditions were not optimal.

Still, transmission could occur via extracellular electrical fields, termed electrical field coupling, and several candidate nanodomains exist for the location of putative ephapses. In 2009, Dr. Delmar began to redefine the understanding of intercalated disc microdomains by discovering the co-distribution of Nav1.5 with structural proteins of the intercalated disc [36], which later led to the discovery of NaV1.5 clusters coordinated by N-cadherin [37, 38]. The foundation for another possible ephapse was laid when Gourdie and his team described the perinexus at the edge of gap junctional plaques [39]. The perinexus represents an extracellular domain where adjacent cardiomyocytes’ membranes separate the gap junctional plaque and other structural junctions such as adherens junctions. Very soon after, Gourdie’s group revealed that Nav1.5 is also co-adjacent to gap junctions in the perinexus [40]. Finally, in collaboration with Gourdie, Smyth, Keener, and Lin, we have repeatedly demonstrated that expanding perinexal width is a common mechanism associated with slowed cardiac conduction in a variety of species, chambers of the heart, and conditions [22, 24, 25, 41-43]. Alternative to electric field coupling, ephaptic mechanisms could also include local depolarization via accumulation of extracellular potassium. Our experiments cannot discern between these possibilities.

The demonstration that ephaptic transmission is possible, does not prove that it actually occurs or that it has any functional significance. Thus, we continued experiments in native pairs that exhibited varying degrees of electrical coupling and AP activation delays that were shorter, in the range of a few msec. This suggests that EtC is a more efficient transmission mechanism; however, several observations suggest that non-gap junctional mechanisms contribute. Although the passive cardiomyocyte depolarized during the stimulus, this occurred with some delay, a delay that only weakly correlated with gap junctional coupling. Even the active cell did not depolarize with a single time constant, but rather a linear depolarization was initiated and the rate maintained upon termination of the stimulus. This behavior is indicative of an active process, which was mirrored by the passive cell. In support of this being an active process rather than an EtC phenomenon, the rate of depolarization in the passive cell did not change upon stimulus termination despite the fact that the driving gradient was eliminated. This suggests that the initial depolarization of the active cell was sufficient to trigger a subset of channels in the passive cell. Furthermore, the current of this subset was large enough to activate further sodium channels to reach the all-or-none activation threshold for initiating an action potential.

A more direct indication that gap junctions were not the sole mediator of transmission was evidenced by the lack of effect of electrical coupling on transmission delay. Thus, even as gap junctional coupling approached zero, this did not alter AP delay between cardiomyocytes. This is in contrast to computational studies incorporating only gap junction mediated conduction, which predict that AP propagation cannot occur with gap junctional coupling below 5.6 nS [44]. Here we show that APs transmit with no additional delay down to 1.14 nS coupling. Furthermore, one of the de novo formed pairs developed 1.39 nS of electrical coupling over time, and although this shortened the activation delay from an initial 16.9 msec to 7.9 msec, it did not approach the maximum 0.3 msec delay observed in the native pairs with a comparable electrical coupling. Importantly, the point is not that gap junctions are unimportant. Rather, this study emphasizes that gap junctional coupling is one mechanism of electrical communication between cardiomyocytes that may operate in parallel to ephaptic communication. In accordance with this, studies show that reducing either gap junctional or ephaptic communication, makes the heart more vulnerable to changes in the other, e.g. [45], see [46] for review.

We also show that activation of sodium current in one cell of a pair always results in the activation of sodium channels in the passive neighbor. This occurred despite a quite negative holding potential and irrespective of the degree of coupling. It may be argued that technical limitations in the ability to adequately clamp the active cell, could allow for local depolarization of the passive cell. It is well known that the geometrical shape of a patched cell alters both the intracellular and extracellular electric fields when currents are passed through the system. Even though cardiomyocytes are relatively small and intra- and extracellular-fields are often considered isopotential, the long-axis of a cardiomyocyte and associated membrane capacitance and resistance is different from the short axis, and therefore not every part of the membrane is held at precisely the holding potential set by the amplifiers. This limitation actually works to the advantage of ephaptic coupling because the phenomenon relies on the inability to control the extracellular potential of the tightly, structurally coupled intercalated disc. Mathematical models actually predict that the presence of isolated domains with ephaptic transmission will induce such behavior [47]. Still, this concern could possibly be addressed by reducing coupling or sodium currents to dampen the voltage drop from pipette to more distant membrane domains. To circumvent the space clamp problem, we performed the steady state inactivation experiments.

A central prerequisite for EpC is the existence of partly “shielded” domain, where local extracellular potentials and ion concentrations can vary relative to the bulk extracellular space and allow for biophysical responses that differ compared to whole-cell patch clamping of isolated single cardiomyocytes. We hypothesized that such a domain should contain all elements required for AP formation. This would include potassium channels to maintain a negative resting potential. Consequently, the shielded domain should be able to repolarize during a long-enforced depolarization of a cardiomyocyte pair. The experiments showed that after holding the cardiomyocyte pairs at potentials that would normally maintain the channels in an inactivated state, a test pulse to 0 mV elicited an inward current. These currents were likely sodium channels that had inactivated during the early phase of the pre-pulse, but which resided in a domain where potassium currents, like I_K1_, [48] had repolarized the local potential and allowed the channels to de-inactivate. This was supported by the lack of current briefly after initiating a pre-pulse to -40 mV, the slow time course of current recovery and the sensitivity to barium. Determining the area under curve above -40 mV, could present itself as a novel metric for the ephaptic population of sodium channels.

As mentioned in the introduction, ephaptic coupling has been supported by mathematical modeling, first by Sperelakis and when it was demonstrated that Nav1.5 is densely expressed in the intercalated disc [2] the interest was reinforced. In 2002, Kucera, Rohr and Rudy recapitulated the finding of dense Nav1.5 localization in the intercalated disc. With a more sophisticated model than originally proposed by Sperelakis, these investigators predicted testable relationships that would support the existence and importance of EpC [12]. In brief, the models predicted that (1) cardiac conduction should be more sensitive to gap junctional uncoupling when intercellular separation between apposing Nav1.5 channels within intercalated discs is wide, and (2) separating cardiomyocytes (reducing EpC) can slow cardiac conduction.

Mori, Fishman and Peskin confirmed these two experimental predictions for EpC and conduction, while also developing a model to explore the importance of ionic concentration fluctuations in the intercalated disc [15]. These predictions were recapitulated by Keener and Lin, while further introducing the concept that intercellular propagation, particularly in the transverse direction, can occur by considering the intercalated disc an inverted cable where the ion channels facing these narrow clefts can propagate the *extracellular* electrical signal between cells [49, 50].

To our knowledge the data presented in the present study, are the first direct evidence that supports the model predictions and the growing number of more indirect experimental evidence of ephaptic communication between cardiomyocytes. We show for the first time that ephaptic transmission is possible, that it is most likely in operation, also in the presence of gap junctional coupling, and finally our study suggests that data obtained in single cells may fail to reproduce key features of the electrophysiology of cardiomyocytes that interact with neighboring cardiomyocytes.

## Notes

### Competing Interest Statement

The authors have declared no competing interest.

## References

1. Nielsen, M.S., et al., The intercalated disc: a unique organelle for electromechanical synchrony in cardiomyocytes. Physiol Rev, 2023. 103(3): p. 2271–2319.

2. Cohen, S.A., Immunocytochemical localization of rH1 sodium channel in adult rat heart atria and ventricle. Presence in terminal intercalated disks. Circulation, 1996. 94(12): p. 3083–6.

3. Lin, X., et al., Subcellular heterogeneity of sodium current properties in adult cardiac ventricular myocytes. Heart Rhythm, 2011. 8(12): p. 1923–30.

4. Weidmann, S., The electrical constants of Purkinje fibres. J Physiol, 1952. 118(3): p. 348–60.

5. Kleber, A.G., C.B. Riegger, and M.J. Janse, Electrical uncoupling and increase of extracellular resistance after induction of ischemia in isolated, arterially perfused rabbit papillary muscle. Circ Res, 1987. 61(2): p. 271–9.

6. Hiramatsu, Y., et al., Rate-dependent effects of hypoxia on internal longitudinal resistance in guinea pig papillary muscles. Circ Res, 1988. 63(5): p. 923–9.

7. Freeman, L.C. and W.W. Muir, 3rd, Effects of halothane on impulse propagation in Purkinje fibers and at Purkinje-muscle junctions: relationship of Vmax to conduction velocity. Anesth Analg, 1991. 72(1): p. 5–10.

8. Xing, D., et al., ZP123 increases gap junctional conductance and prevents reentrant ventricular tachycardia during myocardial ischemia in open chest dogs. J Cardiovasc Electrophysiol, 2003. 14(5): p. 510–20.

9. Haugan, K., et al., The antiarrhythmic peptide analog ZP123 prevents atrial conduction slowing during metabolic stress. J Cardiovasc Electrophysiol, 2005. 16(5): p. 537–45.

10. Picone, J.B., N. Sperelakis, and J.E. Mann, Expanded model of the electric field hypothesis for propagation in cardiac muscle. Mathl Comput Modelling, 1991. 15(8): p. 17–35.

11. Sperelakis, N., An electric field mechanism for transmission of excitation between myocardial cells. Circ Res, 2002. 91(11): p. 985–7.

12. Kucera, J.P., S. Rohr, and Y. Rudy, Localization of sodium channels in intercalated disks modulates cardiac conduction. Circ Res, 2002. 91(12): p. 1176–82.

13. Tsumoto, K., et al., Roles of subcellular Na+ channel distributions in the mechanism of cardiac conduction. Biophys J, 2011. 100(3): p. 554–63.

14. Lin, J. and J.P. Keener, Ephaptic coupling in cardiac myocytes. IEEE Trans Biomed Eng, 2013. 60(2): p. 576–82.

15. Mori, Y., G.I. Fishman, and C.S. Peskin, Ephaptic conduction in a cardiac strand model with 3D electrodiffusion. Proc Natl Acad Sci U S A, 2008. 105(17): p. 6463–8.

16. Poelzing, S., S.H. Weinberg, and J.P. Keener, Initiation and entrainment of multicellular automaticity via diffusion limited extracellular domains. Biophys J, 2021. 120(23): p. 5279–5294.

17. Morris, J.A., et al., Nernst-Planck-Gaussian modelling of electrodiffusional recovery from ephaptic excitation between mammalian cardiomyocytes. Front Physiol, 2023. 14: p. 1280151.

18. Moise, N., et al., Intercalated disk nanoscale structure regulates cardiac conduction. J Gen Physiol, 2021. 153(8).

19. Ivanovic, E. and J.P. Kucera, Tortuous Cardiac Intercalated Discs Modulate Ephaptic Coupling. Cells, 2022. 11(21).

20. Suenson, M., Ephaptic impulse transmission between ventricular myocardial cells in vitro. Acta Physiol Scand, 1984. 120(3): p. 445–55.

21. Veeraraghavan, R., M.E. Salama, and S. Poelzing, Interstitial volume modulates the conduction velocity-gap junction relationship. Am J Physiol Heart Circ Physiol, 2012. 302(1): p. H278–86.

22. Veeraraghavan, R., et al., Sodium channels in the Cx43 gap junction perinexus may constitute a cardiac ephapse: an experimental and modeling study. Pflugers Arch, 2015.

23. Adams, W.P., et al., Extracellular Perinexal Separation Is a Principal Determinant of Cardiac Conduction. Circ Res, 2023. 133(8): p. 658–673.

24. Raisch, T.B., et al., Intercalated Disk Extracellular Nanodomain Expansion in Patients With Atrial Fibrillation. Front Physiol, 2018. 9: p. 398.

25. George, S.A., et al., Modulating cardiac conduction during metabolic ischemia with perfusate sodium and calcium in guinea pig hearts. Am J Physiol Heart Circ Physiol, 2019. 316(4): p. H849–H861.

26. Weingart, R. and P. Maurer, Action potential transfer in cell pairs isolated from adult rat and guinea pig ventricles. Circ Res, 1988. 63(1): p. 72–80.

27. Matteucci, C., Sur un phenomene physiologique produit par les muscles en contraction. Ann Chim Phys, 1842. 6.

28. Katz, B. and O.H. Schmitt, Electric interaction between two adjacent nerve fibres. J Physiol, 1940. 97(4): p. 471–88.

29. Arvanitaki, A., EFFECTS EVOKED IN AN AXON BY THE ACTIVITY OF A CONTIGUOUS ONE. J Neurophysiol, 1942. 5(2): p. 89–108.

30. Sperelakis, N., T. Hoshiko, and R.M. Berne, Nonsyncytial nature of cardiac muscle: membrane resistance of single cells. Am J Physiol, 1960. 198: p. 531–6.

31. Sperelakis, N., Additional evidence for high-resistance intercalated discs in the myocardium. Circ Res, 1963. 12: p. 676–83.

32. Tarr, M. and N. Sperelakis, Weak Electrotonic Interaction between Contiguous Cardiac Cells. Am J Physiol, 1964. 207: p. 691–700.

33. Sperelakis, N., G. Mayer, and R. Macdonald, Velocity of propagation in vertebrate cardiac muscles as functions of tonicity and [K+]. Am J Physiol, 1970. 219(4): p. 952–63.

34. Cole, W.C., J.B. Picone, and N. Sperelakis, Gap junction uncoupling and discontinuous propagation in the heart. A comparison of experimental data with computer simulations. Biophys J, 1988. 53(5): p. 809–18.

35. Sperelakis, N., Combined electric field and gap junctions on propagation of action potentials in cardiac muscle and smooth muscle in PSpice simulation. J Electrocardiol, 2003. 36(4): p. 279–93.

36. Sato, P.Y., et al., Loss of plakophilin-2 expression leads to decreased sodium current and slower conduction velocity in cultured cardiac myocytes. Circ Res, 2009. 105(6): p. 523–6.

37. Agullo-Pascual, E., et al., Super-resolution imaging reveals that loss of the C-terminus of connexin43 limits microtubule plus-end capture and NaV1.5 localization at the intercalated disc. Cardiovasc Res, 2014. 104(2): p. 371–81.

38. Leo-Macias, A., et al., Nanoscale visualization of functional adhesion/excitability nodes at the intercalated disc. Nat Commun, 2016. 7: p. 10342.

39. Rhett, J.M., J. Jourdan, and R.G. Gourdie, Connexin 43 connexon to gap junction transition is regulated by zonula occludens-1. Mol Biol Cell, 2011. 22(9): p. 1516–28.

40. Rhett, J.M., et al., Cx43 associates with Na(v)1.5 in the cardiomyocyte perinexus. J Membr Biol, 2012. 245(7): p. 411–22.

41. George, S.A., et al., Extracellular sodium dependence of the conduction velocity-calcium relationship: evidence of ephaptic self-attenuation. Am J Physiol Heart Circ Physiol, 2016. 310(9): p. H1129–39.

42. George, S.A., et al., TNFalpha Modulates Cardiac Conduction by Altering Electrical Coupling between Myocytes. Front Physiol, 2017. 8: p. 334.

43. Veeraraghavan, R., et al., The adhesion function of the sodium channel beta subunit (beta1) contributes to cardiac action potential propagation. Elife, 2018. 7.

44. Shaw, R.M. and Y. Rudy, Ionic mechanisms of propagation in cardiac tissue. Roles of the sodium and L-type calcium currents during reduced excitability and decreased gap junction coupling. Circ Res, 1997. 81(5): p. 727–41.

45. George, S.A., et al., Extracellular sodium and potassium levels modulate cardiac conduction in mice heterozygous null for the Connexin43 gene. Pflugers Arch, 2015.

46. George, S.A. and S. Poelzing, Cardiac conduction in isolated hearts of genetically modified mice - Connexin43 and salts. Prog Biophys Mol Biol, 2016. 120(1-3): p. 189–98.

47. Hichri, E., H. Abriel, and J.P. Kucera, Distribution of cardiac sodium channels in clusters potentiates ephaptic interactions in the intercalated disc. J Physiol, 2018. 596(4): p. 563–589.

48. Veeraraghavan, R., et al., Potassium channels in the Cx43 gap junction perinexus modulate ephaptic coupling: an experimental and modeling study. Pflugers Arch, 2016. 468(10): p. 1651–61.

49. Copene, E.D. and J.P. Keener, Ephaptic coupling of cardiac cells through the junctional electric potential. J Math Biol, 2008. 57(2): p. 265–84.

50. Lin, J. and J.P. Keener, Modeling electrical activity of myocardial cells incorporating the effects of ephaptic coupling. Proc Natl Acad Sci U S A, 2010. 107(49): p. 20935–40.

